# Structure-guided design of protein attachment points for functional augmentation of complex molecular machines

**DOI:** 10.1101/2025.06.22.660945

**Authors:** Georgie Hau Sørensen, Jessica James, Harrison Steel, Fabio Parmeggiani, Thomas E. Gorochowski

## Abstract

Protein-protein conjugation systems are a powerful way of creating fusion proteins and enable the dynamic combination of protein domains with diverse functionalities. However, the insertion of these systems into enzymes is often performed with little consideration of the structural impact they might have. This is particularly relevant when modifying complex molecular machines that transition between numerous conformational states. Here, we address this issue by developing SIMPLIFE, a computational workflow that supports the design of optimal insertion sites for conjugation tags based on the structure of the proteins involved and performs localised residue redesign where needed. We demonstrate how SIMPLIFE can be used to effectively augment the function of T7 RNA polymerase using the DogCatcher-DogTag system, enabling diverse and dynamically varying mutations within a targeted region of DNA. This work demonstrates the power of combining biophysical and machine learning based approaches for protein structure prediction to efficiently augment the function of molecular machines, accelerating our ability to combine complex biochemical functionalities in new ways.

## INTRODUCTION

Structure-guided protein design using computational tools like Rosetta [1] holds much promise for accelerating the engineering of biology. This approach has already been used for broad applications spanning the design of new-to-nature antibodies with designed specificity [2] and the creation of custom three-dimensional protein structures that could act as scaffolding for synthetic cells or to spatially co-ordinate multi-step biochemical processes [3]. Structure-guided protein design is also particularly well-suited to augmenting the function of existing proteins, extending or combining molecular functions in new ways through the fusion of functional enzymes and domains to blend biomolecular functionalities in ways not typically found in nature [4–8].

Structure-guided protein design is often reliant on experimental protein structures collected using cryogenic electron microscopy (Cryo-EM) or X-ray crystallography, which is expensive to produce [9]. A possible solution to this limitation is to exploit recent advances in computational protein structure prediction. Machine learning based models like AlphaFold [9, 10] are now able to accurately infer the structure of many proteins from amino acid sequence alone. The use of computationally-predicted structures for the rational design of proteins could expand the range of possible designs and is especially valuable when working with proteins that are difficult to crystallise.

When designing new proteins it is often necessary to combine multiple existing biochemical functionalities in novel ways. Recent examples of this are the numerous genome editing technologies that combine the programmability of CRISPR-Cas systems to target specific DNA sequences with other DNA modifying functions for base mutations [11], prime editing [7] and DNA integration [8]. Furthermore, the colocalisation of enzyme domains has also been shown to enable beneficial channelling of metabolites in multi-step pathways [4]. In almost all cases, the combination of such functions is achieved through in-frame translational fusion of entire enzymes or protein domains at their termini, with flexible linker sequences often used to reduce the chance of unwanted steric interference [12].

An alternative approach to the synthesis of a single multi-domain protein, is the use of protein-protein conjugation systems like the SpyTag-SpyCatcher system [13] (**Figure 1**). These enable the dynamic attachment of multiple proteins with differing functions through the formation of an isopeptide bond between a ‘catcher’ protein and a small peptide ‘tag’. The small size of these tags (typically 13–23 amino acids long) means that they are less likely to have a detrimental effects on the structure of the final fusion protein. Furthermore, unlike translational fusions, the use of attachments opens up new opportunities. For example, varying the expression of an attachment can affect the number of sites bound, having multiple types of attachment enables varying stochiometry of additional functional domains, and all these features can potentially vary over time.

**Figure 1:**
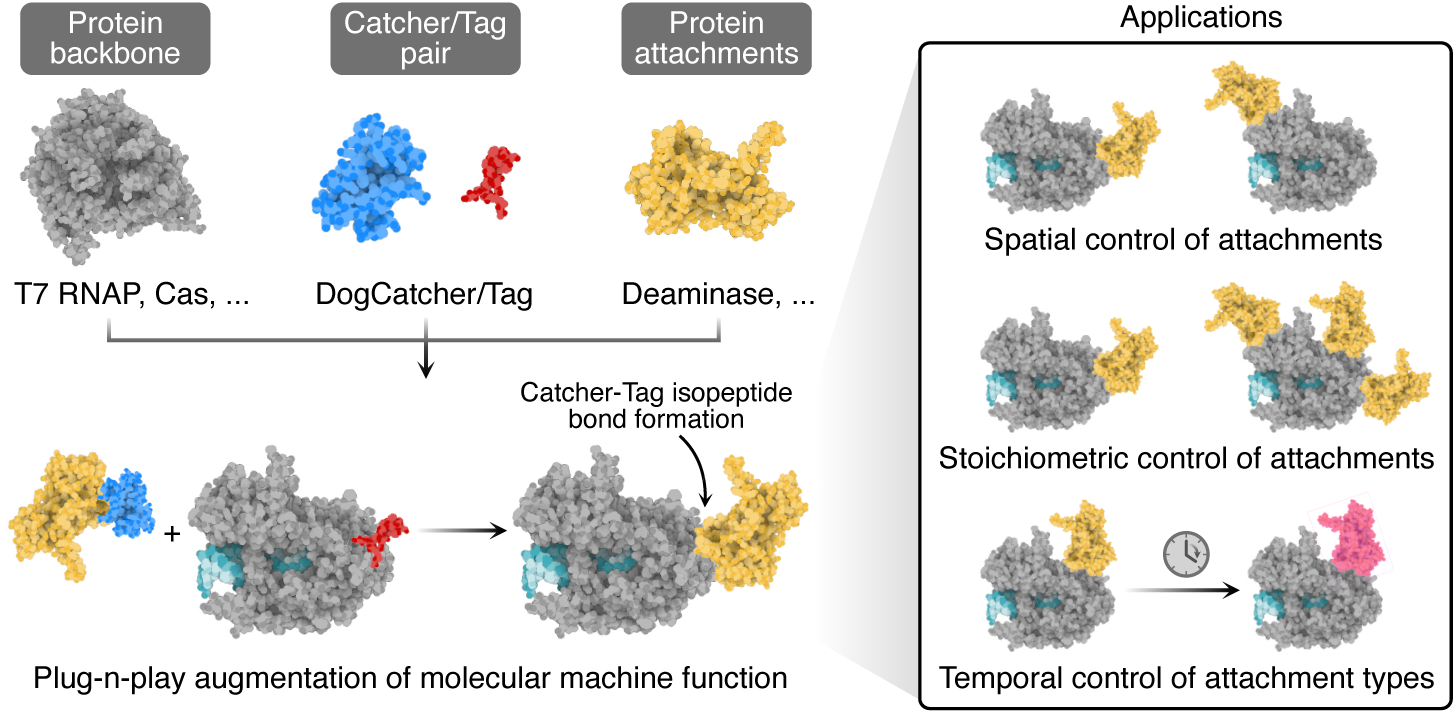
Functional augmentation of molecular machines using protein-protein conjugation tags as attachment points. The use of protein conjugation tags enables control over the spatial positioning of attachments, offers the ability to add multiple sites for stoichiometric control of the component parts, and allow for dynamically varying attachment types.

Most tag-catcher systems developed to date have been designed for use at the termini of proteins, primarily because those points are often accessible and enable efficient bond formation. While offering two possible sites for attachment, this approach limits the overall number of attachments that can be made and the physical positioning of the attached protein. To overcome this limitation, catcher-tag systems like

DogCatcher-DogTag have been developed that are optimised for insertion into protein loops [14]. The applications of protein binding tags have so far mostly ignored the structural implications of their insertion into a catalytically active proteins. However, given the effect even minor changes in loop structure can have on protein function [15, 16], a näıve approach where tags are inserted randomly into any possible loop is likely to diminish activity of the host protein. Some limited studies have explored the potential to use structural modelling when designing insertions [17], but these have so far lacked any functional assessment of their designs and have not made use of accurate AI-based structural models to further refine their suggestions.

In this work, we address this need by developing a new computational workflow called SIMPLIFE (Structurally Informed Modification of Protein Loops for Insertion of Functional Entities) that supports the discovery of optimal insertion points for peptide tags based on the structures of the proteins involved. We demonstrate how SIMPLIFE can be used to functionalise T7 RNA polymerase (T7-RNAP) in several different ways using the DogCatcher-DogTag system. We identify numerous sites for peptide insertion and demonstrate that tags inserted at these locations are more likely to produce engineered polymerases with enhanced activity compared to use of a non-structure-guided approach. Furthermore, we show how the function of these T7-RNAP variants can be dynamically altered via changing expression of the tag component to enable varying types and efficiencies of mutagenesis. This work demonstrates the power of combining biophysical and machine learning based approaches for protein structure prediction to efficiently augment the function of molecular machines and accelerate our ability to combine biochemical functionalities in new ways.

## RESULTS

### Computational design of peptide tag insertions using SIMPLIFE

SIMPLIFE is designed to support the insertion of small protein-binding peptide tags into a larger globular host protein, carrying out checks to help ensure the structure of both insert and protein are maintained and carrying out localised residue redesign if needed. Crucially, it removes the need for any prior experimental knowledge regarding parts of the protein that are robust to modification. This is achieved by combining several bio-physical models from the Rosetta suite and the deep-learning based AlphaFold2 model to create protein structure predictions that combine the strengths of each approach. A user provides the sequence of the protein in which they want to make inserts and the peptide tag that will be used as the attachment point for other proteins. SIMPLIFE then uses this information to carry out three main tasks related to grafting, refinement, and validation of potential designs (**Figure 2a**).

**Figure 2:**
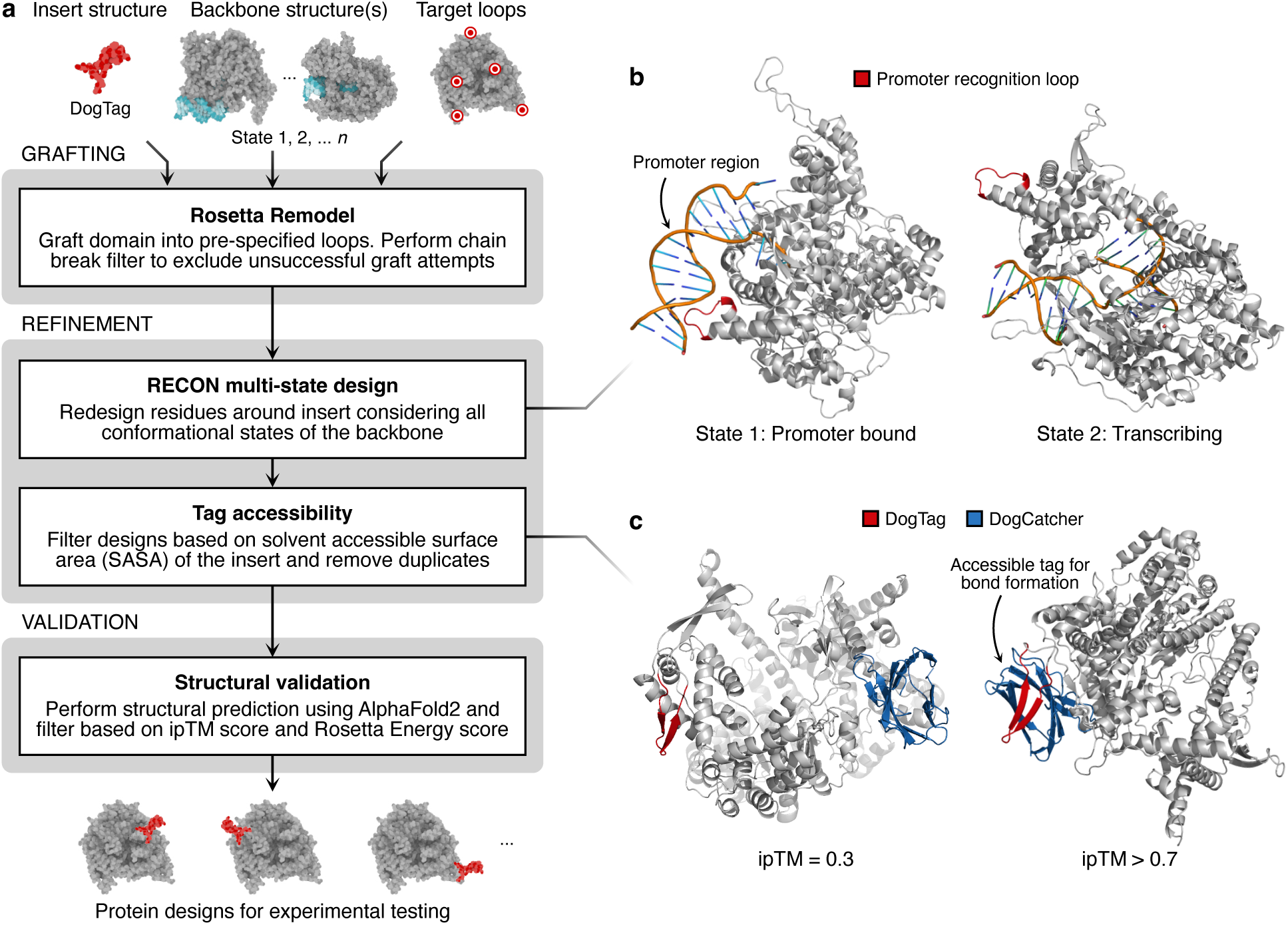
Overview of the SIMPLIFE workflow. (**a**) Diagram of the general structure of the SIMPLIFE workflow, including three core modules (**b**) Protein structure visualisation demonstrating the difference in protein conformation for T7-RNAP when bound to a promoter (PDB: 1H38), or transcribing RNA (PDB: 1CEZ), respectively. (**c**) Visualisation of the protein structures for T7-RNAP::DogTag with bound DogCatcher where the accuracy of the predicted relative positions of DogCatcher and T7-RNAP::DogTag is low (ipTM = 0.3) or high (ipTM *>* 0.7)

In the first step, a rough graft of the peptide structure is made into the target protein using the Rosetta Remodel tool [18]. Remodel attempts to extend each potential protein loop with the sequence of the peptide tag to be inserted. Additional residues on either side of the insert are then selected for subsequent redesign to optimise the insert such that it has minimal impact on the original structures of the host protein and tag. To speed up this process, only the centroid modelling part of Remodel is used for the grafting module; full-atom modelling is performed during the subsequent refinement step.

Once initial grafts have been created, a refinement step then uses the RECON tool [19] to perform a multi-state redesign of residues flanking the inserted peptide. RECON’s multi-state design algorithm allows for consideration of multiple possible conformational states to ensure the inserts will not interfere with protein functions that require dynamic changes in the overall structure (**Figure 2b**). To ensure accessibility of the inserted tag, output from RECON is further filtered based on the surface availability of the inserted tag using a Solvent Accessible Surface Area (SASA) metric. Duplicated sequences are also discarded at this stage.

Finally, a validation step generates structure predictions for each potential design using the AlphaFold Multimer extension of AlphaFold2 [20]. These predictions are filtered based on interface predicted template modelling (ipTM) scores that measure the accuracy of the predicted positions of protein subunits relative to each other (**Figure 2c**). The filter retains predictions with ipTM *>* 0.7, a cut-off chosen based on previously published benchmarking data [21]. Predictions passing this filter are superimposed on their corresponding structure from the RECON module and the resulting root mean squared deviation (RMSD) calculated. A filter of RMSD *<* 5 is applied to remove designs with a large discrepancy between their predicted structure from either AlphaFold2 or RECON.

### T7 RNA polymerase designed for plug-and-play protein attachments

To demonstrate the ability for SIMPLIFE to modify complex proteins with attachment points for later functionalisation, we selected the T7-RNAP and DogCatcher-DogTag system. T7-RNAP is commonly used to transcribe heterologous genes due to its orthogonality to endogenous transcriptional machinery and its high processivity, ensuring strong expression of a desired gene. It also performs a function that requires large conformational changes in its structure over time and therefore acts as a challenging starting point for our workflow.

We used SIMPLIFE to design insertions of the DogTag peptide sequence flanked by `GS` residues into T7-RNAP. We provided a total of 172 possible insert locations across 16 separate loops (*a*–*p*) of the T7-RNAP (**Figure 3a**; **Supplementary Table 1**). These loops were chosen by identifying those present in all known structural conformations of the RNAP with a length ≥ 6 residues. From these possible insert locations, SIMPLIFE then generated 46 unique output designs across 10 different loops. A subset of 20 of these were chosen for experimental characterisation based on how well original structures were maintained and to cover the diversity of the designs produced (**Figure 3a**).

**Figure 3:**
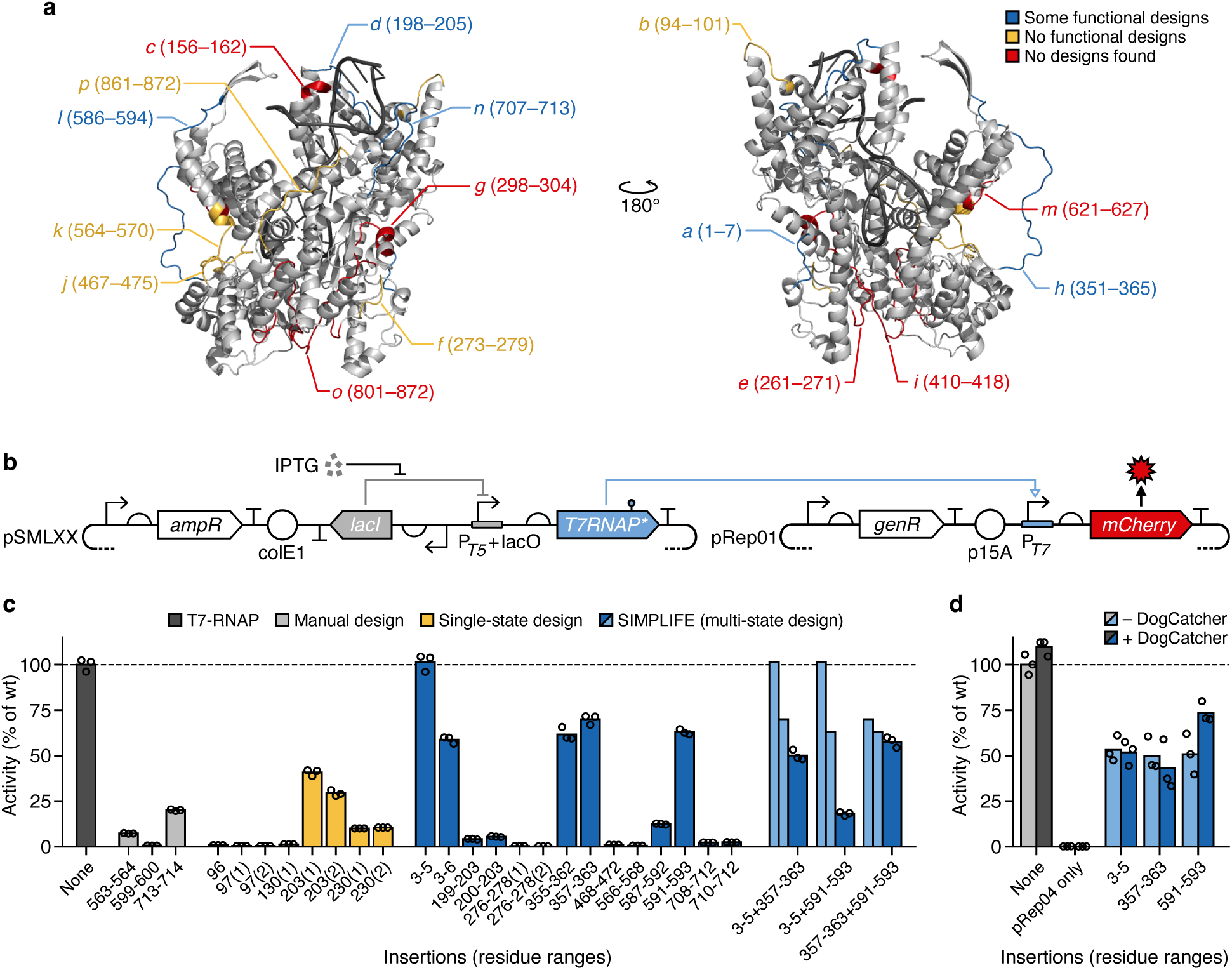
Characterisation of DogTag insertions into T7 RNA polymerase. (**a**) Structural overview of all T7-RNAP loops (*a*–*p*) evaluated by SIMPLIFE (PDB: 1CEZ). Loops are colour-coded by their outcome: red = no designs found; yellow = designs found and some tested experimentally, but none found to be functional; and blue = designs found and some confirmed experimentally to be functional. (**b**) Schematic of the two-plasmid system used for assessing the transcriptional activity of T7-RNAP variants. T7RNAP* denotes the T7-RNAP with one or more DogTag insertions. Genetic diagrams shown using Synthetic Biology Open Language (SBOL) Visual notation [40]. (**c**) Activity of different single and double DogTag insertions into T7-RNAP normalised to the activity of an unmodified T7-RNAP. Bars represent the mean of 3 biological replicates (individual measurements shown by unfilled circles). Thin light blue coloured bars denote the activities of the two single insertions for the associated double insertion (in darker blue) next to it. ‘None’ denotes an unmodified T7-RNAP and dashed line the activity of an unmodified T7-RNAP. (**d**) Activity of high performing single insertions with (dark coloured) and without (ligh coloured) the DogCatcher protein present normalised to the activity of an unmodified T7-RNAP. Bars represent the mean of 3 biological replicates (individual measurements shown by unfilled circles). ‘pRep04 only’ corresponds to cells only containing the pRep04 reporter plasmid. Dashed line denotes the activity of an unmodified T7-RNAP.

High-resolution crystal structures for the T7-RNAP show that the polymerase exists in two different structural conformations corresponding to when it is binding its cognate promoter and carrying out transcription [22]. To test whether the multi-state design employed by SIMPLIFE helped to design better inserts, we also carried out a parallel workflow that omitted any multi-state considerations (**Methods**). We also manually designed several T7-RNAPs for comparison. In this case, we identified 3 different loops through visual inspection of the structure. Sites were chosen based on their location in the middle of large loops (*>*6 residues) that exists on the outer surface of the protein.

To assess whether the modified T7-RNAPs retained their ability to transcribe DNA, we carried out an expression assay using a plasmid containing a fluorescent mCherry reporter protein driven by a T7 promoter (**Figure 3b**). Cells co-transformed with both this plasmid (pRep01) and a plasmid enabling inducible expression of a modified T7-RNAP (pSML01–pSML27) were grown for 16 hours in a plate reader. We found that the activity of the T7-RNAP variants, in terms of fluorescence per cell, varied during different growth phases (**Supplementary** Figure 1). Therefore, for comparisons we considered the the maximum fluorescence per cell during exponential growth (**Methods**) and normalised this activity to the activity of an unmodified T7-RNAP to assess the impact of the insertions on function.

Analysis of our data showed that most insertions caused a large drop in T7-RNAP function (**Figure 3c**). This was particularly prominent for manual designs and those designed using the workflow that omitted multi-state considerations. For designs generated by both these approaches only 2 out of the 11 modified T7-RNAPs maintained *>*25% of their original activity and 5 of the designs showed a complete loss of transcription. In contrast, 5 out of 14 designs generated by SIMPLIFE maintained *>*50% activity, approximately doubling the success rate and average activity achieved. The best performing SIMPLIFE sites were near the N-terminal of the T7-RNAP (residues 3–5), displaying activity similar to the native T7-RNAP, and sites near the loops *h* (residues 351–365) and *l* (residues 586–594), which retained 74% and 63% of wild-type activity, respectively. Interestingly, insertions around the loop near residue 203 saw better activity for the non-multi-state design approach (**Figure 3c**). This likely stems from slight differences in the way that the inserts are grafted into the backbone and highlights the sensitivity that some proteins can have to even small modifications at critical points in their structure.

To test the ability for SIMPLIFE to carry out more extensive modifications, we allowed for two insertions to be made within the same T7-RNAP backbone. These designs were constructed by selecting previous insertions that alone allowed the T7-RNAP to retain good activity. For loops with multiple inserts, a single representative site was chosen. This resulted in three T7-RNAPs combining pairs of insertions at residues 3–5, 357–363 and 591–593.

Activities of the resultant multi-insert T7-RNAPs were again characterised via the same expression assay using plasmids that enables inducible expression of the modified T7-RNAPs (pSML028–pSML030). We found that all of the multi-insert designs displayed reduced activity compared to the single insertions they were constructed from (**Figure 3c**). However, the drops in activity were not additive in nature, with some multiple insertions, like at residues 357-363 and 591-593, showing only small reductions of *<*9% compared to the worst activity of the single insertions. This suggests that even more substantial modification of the T7-RNAP may be possible with only minor impacts on function. However, finding sites that do not synergistically degrade function remains a challenge if large experimental screens are not feasible.

Finally, we carried out experiments with the three best performing single insertions to assess whether co-expression of a DogCatcher protein might lead to changes in T7-RNAP activity due to bond formation with the inserted DogTag. This was tested using a modified reporter plasmid that included an inducible expression cassette for the DogCatcher protein (pRep04). T7-RNAP activity was then measured with and without co-expression of the DogCatcher protein.

We found slightly lower activities for all of the insertions tested and greater variation in the activities measured across the biological replicates (**Figure 3d**). This may be due to the additional burden that expression of the DogCatcher protein places on the cell and the larger reporter plasmid present. Insertions at residues 3–5 and 357–363 saw virtually no change in activity when the DogCatcher was expressed. However, the insertion at residues 591–593, showed a notable increase in activity when the DogCatcher protein was present. This could be due to DogCatcher causing beneficial changes to the T7-RNAP structure once bound, synergistically affecting its function. However, it should be noted that the effect is small given the large variability we see across biological replicates.

### A tunable T7-RNAP mutagenesis platform via a deaminase protein attachment

Having created several active T7-RNAP variants with DogTag attachment points, we next explored ways in which we could augment their function. We chose to focus on the attachment of deaminase domains due to their previous successful fusion with T7-RNAPs to create site-directed mutagenesis systems (e.g., eMutaT7) [5, 23]. In these systems, a T7-RNAP is fused at its C-terminal to a deaminase domain such that transcription by the fusion protein induces deamination of specific bases in the DNA being transcribed. This subsequently causes C → T or A → G mutations after DNA replication for cytosine and adenosine deaminases, respectively. By flanking a region to target for mutagenesis with a T7 promoter and terminator, transcription by the T7-RNAP::deaminase fusion in that specific region leads to localisation of the deaminase domain and ultimately increased mutation rates.

To demonstrate that attachment of deaminase domains through DogCatcher-DogTag conjugation would result in an augmented T7-RNAP system able to perform targeted mutagenesis, we employed the same highly active cytosine deaminase protein, PmCDA1, developed for the eMutaT7 system [5]. We translationally fused PmCDA1 to the C-terminal end of the DogCatcher protein using a flexible `G4TG4S` linker, and expressed this fusion protein from a new reporter plasmid (pRep03) using the P_cymRC_ cuminic acid inducible promoter (**Figure 4a**). This allowed us to test the effect of the T7-RNAP and attachment protein expressed in isolation, as well as both the T7-RNAP and attachment protein expressed together.

**Figure 4:**
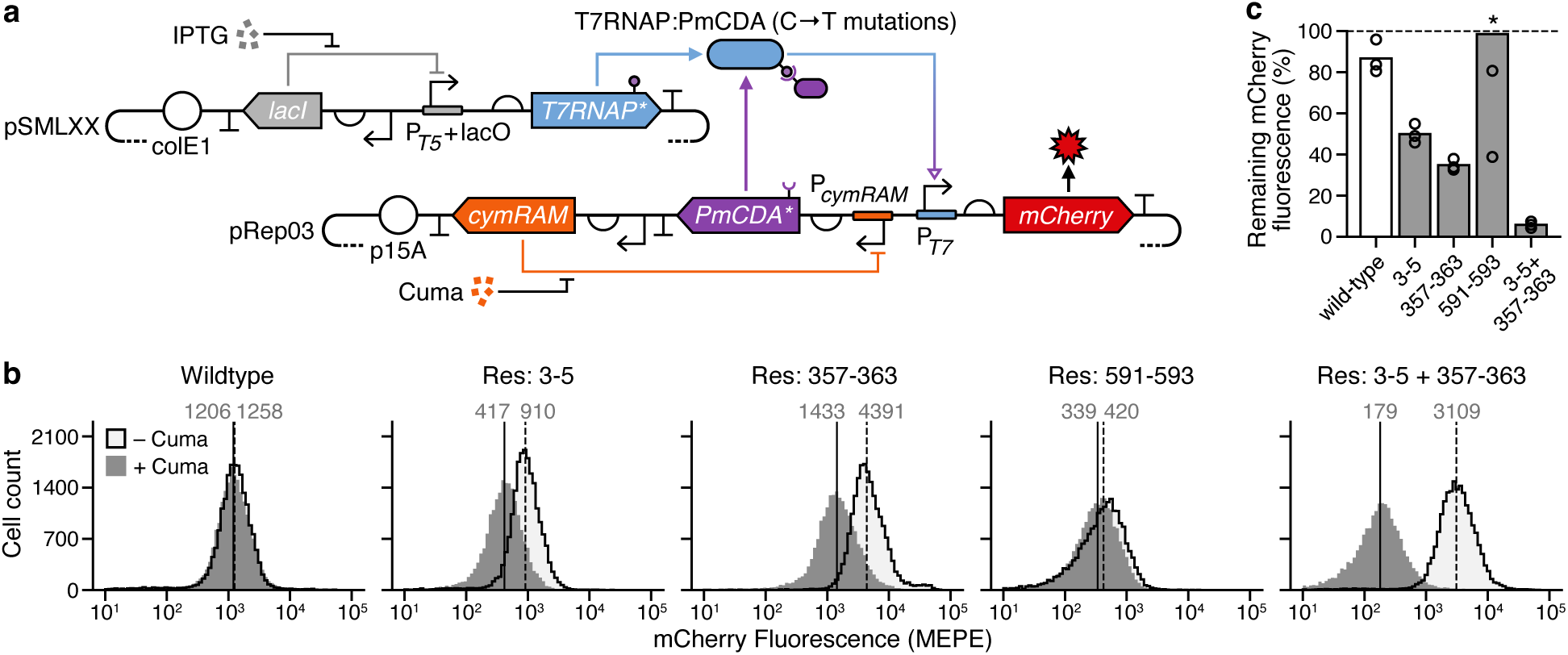
Tagged T7-RNAPs as a tunable targeted mutagenesis platform. (**a**) Schematic of the two plasmid system used for mutagenesis experiments. The pSMLXX plasmid encodes the tagged T7-RNAP containing a single or multiple DogTag insertions (T7RNAP*). The pRep03 plasmid encodes a fluorescence protein reporter used as output (mCherry) and a cuminic acid (Cuma) inducible cytosine deaminase fused to DogCatcher (PmCDA*) to enable localisation to our modified T7-RNAPs. Genetic diagrams shown using Synthetic Biology Open Language (SBOL) Visual notation [40]. (**b**) Representative fluorescence distributions from flow cytometery for different T7-RNAP variants after 32 hours, with (solid grey) and without induction (light grey and solid black outline) by cuminic acid. Solid and dashed vertical lines denote the medians for each distribution with cuminic acid present and absent, respectively, with there median values shown above each line in grey. Fluorescence measurements given in Molecules of Equivalent PE per cell (MEPE) units. For the variant with an insertion at residues 591–593 each distribution varied significantly with drops and increases in fluorescence seen between induced and uninduced conditions (see **Supplementary** Figure 2). (**c**) Remaining mCherry fluorescence for each T7-RNAP variantafter induction of the PmCDA::DogCatcher cytosine deaminase compared to when attachment expression is uninduced. Bars show mean values for 3 biological replicates. Individual values of replicates shown by unfilled circles. The ‘*’ for the variant with insertion at residues 591–593 denotes that one of the value is not shown at 176%.

To characterise the ability of our PmCDA::Dogcatcher attachment to mutate target DNA, we designed a mutagenesis experiment based on the reduction of fluorescence signal from an mCherry reporter protein as detrimental mutations are incorporated into its sequence. Strains transformed with pRep03 and plasmids containing each of the high-performing T7-RNAP variants from the previous characterisation experiments were cultured overnight with T7-RNAP expression induced. This led to baseline production of mCherry when no mutational attachment was present. These cultures were then diluted into fresh media and each split into two groups that were either supplemented with cuminic acid to induce expression of the Dogcatcher::PmCDA1 protein, or not. These cultures were grown for 32 hours with several dilution steps into fresh media (**Methods**), and then samples taken for analysis by flow cytometry to assess shifts in the distribution of fluorescence across the populations.

The vast majority of samples showed a single approximately log-normal distribution of fluorescence. However, as expected, we saw measurable drops in median fluorescence for samples where the mutational attachment was expressed in tandem with a modified T7-RNAP containing an attachment point (**Figure 4b,c**). We found that T7-RNAPs with only single insertions had the smallest drops of 2–50%, although most of these were much larger than the small variation in T7-RNAP activity (∼13%) observed between the absence and presence of the mutational attachment (**Figure 3d**). The differences we observe in the mutational efficiency of the T7-RNAP variants could stem from differences in the physical positioning of the mutational attachments in relation to DNA being transcribed, or may be caused by varying efficiency in the attachment of the mutational domain due to accessibility of the DogTag in the T7-RNAP backbone. However, analysis of the ipTM values (a proxy for tag accessibility to a possible catcher attachment) for the single insertions tested did not show a clear correlation with the drops in fluorescence. There was also no correlation between the transcriptional activity of the modified T7-RNAPs (**Figure 3c**) and the size of the drops in fluorescence. This highlights an important aspect of functional augmentation, where the ability for the attachment to carry out its function is tied to its spatial positioning to other elements within the system. In this case, for the deaminase to function correctly, it will need to interact with the DNA being transcribed, which for some attachment points may be difficult due to occlusion by the T7-RNAP itself. Interestingly, the T7-RNAP multiple-insertions variant, with DogTags at residues 3–5 and 357–363, showed much greater drops in fluorescence, falling 94% (*>*16-fold drop). This likely stems from the increased chance of modification due to a higher density of attachments localised to the target DNA.

These results demonstrate that our engineered T7-RNAPs can be functionalised to varying extents and offer the ability to tune mutational activity, enabling dynamically time varying mutations of target DNA independent of it’s transcription by the same T7-RNAP.

## DISCUSSION

In this work, we have shown how our computational workflow called SIMPLIFE can be used to successfully identify diverse sites for inserting a peptide-binding tag into a large globular protein to enable functional augmentation of the host protein. We have shown that our designed insertions are able to better maintain activity of the host protein compared to manual designs made through visual inspection. This seems to, in part, be a result of the multi-state design approach used by SIMPLIFE, which is able to not only find permissible sites based on multiple possible conformations of the protein, but also to carry out localised redesign of residues to help minimise the impact the insertion has on the host protein’s native set of conformations. However, we have also found one case where the sequence of the designed linkers played a less important role, i.e., insertions made into loop *h* (**Figure 3a**) all resulted in the T7-RNAPs with high transcriptional activity, even though their linker designs differed.

Previous efforts have successfully split T7-RNAP in order to insert light-activated dimerization domains to enable light-based control of transcription [6]. This work identified several previously uncharacterised sites where insertions of the dimerization domains resulted in an active T7-RNAPs. Their best-performing site was between residue residue 563 and 564. Here, we tested insertions in this supposedly permissible location, both through the selection of a SIMPLIFE designed output targetting this loop, and a manually designed insertion (**Figure 3**). In both cases, our modifications resulted in low activity of the engineered T7-RNAP. This demonstrates that permissiveness towards an insertion of one type of domain is not necessarily a good predictor of the permissiveness to other types of domain, even at the same location.

As with all model-based approaches, the quality of the outputs from our workflow depends on the accuracy of initial input structures for both the insert peptide and globular host protein into which insertions are to be made. Here, the interaction of DNA with T7-RNAP is crucial to function. We were able to account for this feature when modelling insertions because of the existence of high-quality crystallography data of T7-RNAP bound to DNA in various key conformations (PDB:1h38 and PDB:1cez). Input structures can be prepared for SIMPLIFE by simply using protein structure prediction tools like AlphaFold2. Unfortunately,

AlphaFold2 did not provide viable protein-DNA models that could have been used as input for SIMPLIFE, and other prediction tools that are able to model protein-nucleotide complexes typically lack the accuracy we need [24]. However, with the release of new foundation models, like AlphaFold3 [25], it is now possible to generate high-quality protein-DNA structures, which are ideal for our workflow, alleviating the need for costly and often challenging structural experiments and opening up its application to less-characterised proteins.

Related to these advances in machine learning, the emergence of diffusion-based models for protein design has also begun to generate traction [26, 27]. These models are typically more computationally expensive than the Rosetta-based biophysical modelling we currently use for our initial grafting and refinement steps, but can be faster in some instances where sufficient computational resources are present. To accommodate such developments, the modular structure of our workflow allows for specific tools at a particular stage to be simply swapped (e.g., allowing us to use a diffusion model for the remodelling step). This ability also opens avenues to workflows tailored for the specific types of protein component being used (e.g., rigid repeat proteins [28]).

In addition to model-based design, high-throughput transposon-based experimental approaches that generate large and diverse insertion libraries for screening offer a complementary approach to finding suitable insertions when activity of a host protein can be easily monitored. However, it should be stressed that such an approach is not capable of finding the designs presented in this work, as transposon-based methods rely on the copy and paste of an identical insert and are unable to redesign nearby residues to accommodate changes the insertion might induce in the host protein’s core structure. We see this ability as key to the success of our approach as most of our best performing insertions do not share similar linker sequences.

The focus of this work was ensuring that newly inserted attachment points minimally affected the host proteins original activity. As highlighted earlier, we did not see any correlation between the activity of the modified T7-RNAPs and the efficiency of mutation (**Figure 3c**), supporting the idea that positioning of the attachment protein is also crucial for its function (i.e., the deaminase coming sufficiently close contact to the transcribed DNA). An interesting future direction, would be to carrying out further filtering of designs based on the spatial positioning of attachments with the goal of enhancing new functionalities, or designing attachments and linkers with more specific geometries to ensure proper positioning of all functional domains [29].

The fact that it is possible to use multiple SIMPLIFE-designed insertions simultaneously within the same protein offers the potential to multi-functionalise a single core protein backbone. Furthermore, our set of internal DogTags for T7-RNAP could also be used in conjunction with other orthogonal catcher/tag systems at the N- and C-termini of T7-RNAP to further functionalise T7-RNAP in a controlled way. Each of the insertion sites in this study has only been computed by SIMPLIFE as single inserts. An interesting future direction would be to use SIMPLIFE to model more than one insertion site simultaneously. This is supported by the workflow, and could result in a better optimised modelling of multi-insert proteins whose activity is less impacted.

The mutagenesis system we develop may also provide an interesting foundation for creating engineered biosystems that are designed to evolve in specific ways. For example, the ability to control the amount and type of mutations that occur [30, 31], e.g., using targetted mutagenesis systems [5, 32], could offer ways to guide evolution and develop more open-ended biological design workflows [33]. The flexibility of our system to having the number and type of attachments varied over time makes it particularly appealing for the control that would be needed for this use case.

The ability to combine biological functions through protein-protein conjugation offers a powerful and complementary approach to designing monolithic molecular machines *de novo*, allowing for multiple proven functionalities to be integrated in new ways that potentially change over time. As synthetic biology moves towards more substantial modifications of biology, this work will support the development of diverse phenotypes required for more complex real-world applications.

## METHODS

### SIMPLIFE workflow

The SIMPLIFE workflow consists of several shell scripts, XML files and Python scripts. Modelling with Rosetta is exclusively done through Rosetta-scripts [34]. To run SIMPLIFE, high quality structure files (i.e., those without gaps or regions of high uncertainty) are required in PDB format for the insert as well as for any known conformational state of the globular target protein. Furthermore, the user must supply a target specification file consisting of two columns where each row denotes an insertion for the SIMPLIFE workflow to model, and the numbers in the two columns representing the start and end residue number of the target protein sequence where the insertion should be attempted. Any residues within the interval given by these start and end specifications will be deleted and replaced by the insertion with SIMPLIFE. Full details of how the workflow is run can be found in our code development repository (**Data Availability**).

### Strains, media and reagents

*Escherichia coli* strain DH10-beta (Δ(ara-leu) 7697 araD139 fhuA ΔlacX74 galK16 galE15 e14-*φ* 80dlacZΔM15 recA1 relA1 endA1 nupG rpsL (StrR) rph spoT1 Δ(mrr-hsdRMS-mcrBC)) (C3019I, New England Biolabs) was used for cloning and propagation of plasmids. Characterisation of T7-RNAP variants was carried out using *E. coli* strain BL21 (F-*dcm ompT hsdS*(rB-mB-) *gal*) (C2530H, New England Biolabs). Cells were grown in lysogeny broth (LB) (L3522, Sigma–Aldrich) for regular outgrowth and propagation. For plate reader experiments to characterise expression of the fluorescent mCherry reporter, as well as mutagenesis experiments, M9 minimal media supplemented with glucose (6.78 g/L Na2HPO4, 3 g/L KH_2_PO_4_, 1 g/L NH_4_Cl, 0.5 g/L NaCl (M6030, Sigma–Aldrich), 0.34 g/L thiamine hydrochloride (T4625, Sigma–Aldrich), 0.4% D-glucose (G7528, Sigma–Aldrich), 0.2% casamino acids (AC61204-5000, Acros), 2 mM MgSO_4_ (213115000, Acros), and 0.1 mM CaCl_2_ (C8106, Sigma–Aldrich) was used. Expression of T7-RNAP variants was controlled by a T5-lacO promoter induced with varying concentrations of Isopropyl-*β* -D-thiogalactopyranoside (IPTG) (R0392, Thermo Fisher Scientific). Expression of DogCatcher fusions to PmCDA1 was induced with 100 µg/mL cuminic acid (268402, Sigma Aldrich), while Dogcatcher fusions to TadA1 were induced with 10 µg/mL anhydrotetracycline hydrochloride (aTc) (233131000, Acros organics). Antibiotic selection of plasmids was performed using either 100 µg/mL ampicillin (Sigma–Aldrich, A9518), 100 µg/mL kanamycin (K1637, Sigma–Aldrich), or 10 µg/mL gentamycin (G3632, Sigma–Aldrich).

### Assessment of tag insertion sites for T7-RNAP

SIMPLIFE was run using the Oracle Cloud Infrastructure (OCI). The total runtime for each successful structure on the cluster was around 5 min. AlphaFold2 was run through Colabfold [10] via an API to execute computations via GPU nodes within the cluster. Input structures were generated as follows: AlphaFold2 was used to generate a PDB file for the DogTag peptide. Two high-quality T7-RNAP structures were obtained from publicly available crystal structures in the PDB database (1H38 and 1CEZ), which represent the two known conformational states of T7-RNAP. For both these structures, small gaps of residues not captured by the X-ray diffraction experiments appeared in the PDB files. Rosetta Remodel was used to fill in these short gaps based on the known protein sequence for T7-RNAP. All input structures were relaxed using Rosetta FastRelax with Cartesian minimization, and a final validation of input structures was performed using MolProbity version 4.5.2 [35] to score a range of fundamental protein geometry variables such as bond angles, bond lengths, and Ramachandran outliers, using the “Analyze geometry without all-atom contacts” function with default settings.

### Construction of T7-RNAP variants

Modified T7-RNAP variants were created using NEBuilder HiFi DNA assembly. The pQE-T7-RNAP plasmid [36], which contains a T7-RNAP coding sequence under the control of a T5-lacO promoter, was used as a starting template. Primers with 5’ overhangs that were complementary to ∼30 bp of the insert amplicon were then used to create specific pQE-T7-RNAP amplicons for each insertion to be made. Primers were manually optimised to avoid primer dimers as the linkers flanking each inserted tag often contained repeats of glycine, which increased the likelihood of dimers forming. Insert amplicons were similarly created as primers containing 5’ overhangs that were complimentary to the backbone amplicon. DNA encoding the DogTag peptide sequence (IPATYEFTDGKHYITNEPI) was synthesized as a gBlock (Integrated DNA Technology). Linkers flanking the insert (5 residues at N-terminal end, and 6 residues at the C-terminal end of tag), each bespoke to the exact insert location as designed by SIMPLIFE, were added through 5’ primer overhangs during amplification. The creation of amplicons for cloning was done by PCR using the Q5 HighFidelity DNA Polymerase (M0491S, New England Biolabs). Following gel extraction using the Monarch

DNA Gel Extraction Kit (T1020S, New England Biolabs), purified amplicons were mixed in a 5:1 molar ratio of insert to backbone, and 4 µL NEBuilder HiFi DNA Assembly Master Mix (E2621, New England Biolabs) was added along with nuclease-free water for a final volume of 8 µL. Following incubation at 50°C for 15 min, 2 µL of each cloning reaction was used to transform 50 µL chemically competent *E. coli* DH10-beta, according to manufacturers protocol. Plates containing the transformed cells were grown overnight at 37°C and individual colonies were then screened by colony PCR using Quick-Load Taq 2X Master Mix (M0271L, New England Biolabs) to confirm the expected insert size. Positive colonies were inoculated into LB liquid cultures and grown overnight at 37°C. Plasmids were extracted the following day using the Monarch Plasmid Miniprep Kit (T1010L, New England Biolabs). To further verify the correct assembly of each T7-RNAP variant plasmid, Sanger sequencing (Source Bioscience) was used, and only sequenceverified plasmids were carried forward for experiments. Details of all T7-RNAP expression plasmids are provided in **Supplementary Table 2**.

### Assembly of reporter plasmids

Reporter plasmids were constructed using NEBuilder HiFi DNA Assembly in a similar way as described above. The initial mCherry reporter plasmid (pRep01) was constructed using PCR fragments from the pSEVA661 plasmid backbone, a synthesised DNA sequence (gBlock, Integrated DNA Technology) encoding a T7 promoter with wild-type Shine-Dalgarno sequence, and PCR fragments of an mCherry coding sequence from the pAB50 plasmid which was a gift from Mustafa Khammash (Addgene plasmid #101678) [6]. These three fragments were combined in equimolar amounts and *E. coli* DH10-beta cells transformed with the assembly mix. The pRep02 plasmid was constructed by using a synthesised gBlock encoding a DogCatcher CDS with in-frame SapI cloning site, flanked by BsaI overhangs (AATG at 5’ end and AGGT at 3’ end of DogCatcher sequence). Using the pCymRAM-sfGF plasmid, a gift from Shivang Joshi, as a template for PCR, fragments for the plasmid backbone were generated, with flanking BsaI overhangs matching the overhangs for the DogCatcher CDS. Golden Gate cloning reaction were subsequently done using BsaI (R3733S, New England Biolabs) and T4 ligase (M0202S, New England Biolabs) to combine the two fragments to form pRepo02. The pRep04 plasmid, containing both mCherry and Dogcatcher, was constructed by PCR amplifying pRep01 to create a backbone fragment into which a PCR fragment containing the PcymRC promoter, DogCatcher CDS, and the cymRAM regulator was cloned by similar methodology to pRep01, using primers with complementary overhangs and NEBuilder HiFi DNA Assembly. The pRep03 plasmid was constructed from the pRep04 plasmid by using the SapI cloning site to insert a synthesised DNA sequence encoding PmCDA1 as used by [5] through Golden Gate cloning. Using overhangs CAG for the 5’end and TAA for the 3’ end of the PmCDA1 sequence, a scarless assembly was done using SapI (R0569S, New England Biolabs) and T4 ligase (M0202S, New England Biolabs).

For all reporter plasmids, cells transformed with cloning reactions were plated on solid LB agar media and subsequent screening of successful colonies was performed similarly to the T7-RNAP assembly approach. Sanger sequencing was used to verify the sequence of the mCherry and T7 promoter in the pRep01 plasmid, the PCymRC promoter and DogCatcher CDS for pRepo02, as well as mCherry and DogCatcher/DogCatcher::PmCDA1 for the pRepo04 and pRepo03 plasmids, respectively.

Details of all reporter plasmids are provided in **Supplementary Table 2**.

### Fluorescence plate reader experiments

Single colonies co-transformed with the mCherry reporter plasmid (pRep01) and each of the respective T7-RNAP variant plasmids (pSML01–pSML30) were used to inoculate 3 mL M9-media with antibiotics in a 24-well deep well plate and incubated at 30°C and 1000 rpm for 16 hours in a shaking incubator (S1505, Stuart). Overnight cultures were subsequently used to inoculate 1 mL pre-cultures by diluting samples to starting OD600 of 0.05 in fresh M9-media supplemented with antibiotics. Following incubation at 30°C and 1000 rpm for an hour in the same shaking incubator, IPTG was added to each culture for a final concentration of 1 mM to induce of T7-RNAP expression. The 1 mL pre-culture was aliquoted into 4 × 200 µL replicates in a black polystyrene clear-bottom 96-well plate (CLS3904, Corning Inc.). Plate reader measurements were then taken using a BioTeK Synergy Neo2 plate reader at 30°C for 16 hours with double orbital shaking. To estimate cell density, OD600 was measured at 15 minute intervals throughout the experiment. Fluorescence from the mCherry reporter was monitored using the instruments monochromator to specify excitation at 560/20 nm, and emission at 610/20 nm, with gain settings of 80 and 120 for each well.

### Mutation screening using flow cytometry

Samples for mutagenesis screening were prepared by inoculating single colonies transformed with pRep03 and varying T7-RNAP variant plasmids into 3 mL M9-glucose media supplemented with kanamycin and gentamycin for selection and 25 µM for induction of T7-RNAP production. Cultures were incubated overnight for 14.5 hrs in a shaking incubator (S1505, Stuart) at 30°C and 1000 rpm using 24 well deep well plates. Overnight cultures were then used to inoculate a new set of 3 mL deep well plate cultures containing M9-glucose plus supplements to a starting OD600 of 0.1. These new inoculated 24-well plates were then placed back into the same shaking incubator at 30°C and 1000 rpm. After 1 hr growth of the cultures, expression of Dogcatcher::PmCDA1 was initiated by adding cuminic acid to a final concentration of 100 µM to each well. Every 8 hours following cuminic acid induction, cultures were diluted into fresh M9-glucose plus supplements in new 24 well plates, with each culture being diluted down to an OD600 of 0.1. After 24 hours, samples for flow cytometry were collected by pipetting 20 µL of each culture into 180 µL PBS with 2 mg/mL kanamycin to halt further gene expression. Remaining volume of mutagenesis culture was miniprepped using the Monarch Spin Plasmid Miniprep Kit (T1110S, New England Biolabs), and the resulting 10 µL eluate in nuclease-free water was saved at –20°C for future analysis.

Flow cytometry samples were kept at room temperature for 1 hr to ensure full maturation of all fluorescent proteins before flow cytometry was performed. Single cell fluorescence of mCherry was then measured using a BD LSRFortessa X-20 Cell Analyzer, using filter settings for the PE fluorochrome (excitation by laser at 488 nm, emission detected at 575/26 nm). Sample collection was capped at a maximum of 100,000 events per sample or an upper time limit of 1 minute per sample. To allow for conversion of mCherry fluorescence into standardised MEFL units, 15 µL 8-peak Rainbow Calibration Particles (RCP-30-5A, Spherotech) was added into a single well containing 200 µL of PBS and calibration/conversions performed using FlowCal version 1.3.0 [37].

### Data analysis and computational tools

Plate reader data was exported as plain CSV files, and analysed using Microsoft Excel. Plots were generated with Python version 3.12.9 using the matplotlib version 3.10, csv version 1.0, FlowCal version 1.3.1, and numpy version 2.0.1 packages. All figures were composed in Affinity Designer version 2.6.3 using Synthetic Biology Open Language (SBOL) Visual glyphs taken from paraSBOLv version 1.0.0 [38]. Molecular visualisations made using PyMol version 3.1.5, and Protein Imager [39].

## DATA AVAILABILITY

Annotated plasmid maps of all constructs used in this study are provided in GenBank format as **Supplementary Data 1**. Code for the SIMPLIFE workflow can be found at: https://github.com/ `BiocomputeLab/SIMPLIFE`.

## ACKNOWLEDGEMENTS

This research was supported by an EPSRC PhD Studentship (G.H.S.), the Oracle for Research program, a UKRI National Engineering Biology Programme Breakthrough Award grant BB/W012448/1 (T.E.G.), the EPSRC-funded EEBio grant EP/Y014073/1 (T.E.G., H.S.), and BrisEngBio, a UKRI-funded Engineering Biology Research Centre grant BB/W013959/1 (T.E.G., F.P.) In addition, T.E.G. was supported by a Turing Fellowship from The Alan Turing Institute under EPSRC grant EP/N510129/1 and a Royal Society University Research Fellowship grant URF/R/221008. F.P. was also supported by a EPSRC early career fellowship EP/S017542/2. The funders had no role in study design, data collection and analysis, decision to publish or preparation of the manuscript.

## AUTHOR CONTRIBUTIONS

T.E.G. conceived the study, secured funding, and performed the sequencing analysis. G.H.S. performed all experiments and analysed the data. J.J. aided with experiments. T.E.G., F.P. and H.S. supervised the work. All authors helped to design experiments, contributed to the interpretation of results, and the writing and editing of the manuscript.

