## Supplementary Information for "Structure-guided design of protein attachment points for functional augmentation of complex molecular machines"

### Supplementary Table 1: Loops in T7 RNA polymerase where modifications made

| Loop | Start residue | End residue | Number of SIMPLIFE output designs | Experimentally tested designs (specific residues) |
|---|---|---|---|---|
| *a* | 1 | 7 | 6 | 3–5, 3–6 |
| *b* | 94 | 101 | 7 | 94–96, 97–100 |
| *c* | 156 | 162 | None | None |
| *d* | 198 | 205 | 3 | 199–203, 200–203 |
| *e* | 261 | 271 | None | None |
| *f* | 273 | 279 | 3 | 276–278, 276–278(2) |
| *g* | 298 | 304 | None | None |
| *h* | 351 | 365 | 6 | 355–362, 357–363 |
| *i* | 410 | 418 | None | None |
| *j* | 467 | 475 | 3 | 468–472, 468–472(2) |
| *k* | 564 | 570 | 4 | 565–568, 565–568(2) |
| *l* | 586 | 594 | 7 | 587–592, 591–593 |
| *m* | 621 | 627 | None | None |
| *n* | 707 | 713 | 6 | 708–712, 710–712 |
| *o* | 801 | 807 | None | None |
| *p* | 861 | 872 | 22 | 862–868, 867–971 |

#### Supplementary Table 2: Plasmids used in this study

| Name | Description | Resistance | Origin |
|---|---|---|---|
| pSML01 | Wild-type T7-RNAP (derived from pQE-WT) | Kanamycin | pBR322 |
| **T7-RNAP expression plasmids with single-state designed insertions** | | | |
| pSML02 | T7-RNAP with DogTag insertion at residues 3-5 | Kanamycin | pBR322 |
| pSML03 | T7-RNAP with DogTag insertion at residues 3-6 | Kanamycin | pBR322 |
| pSML04 | T7-RNAP with DogTag insertion at residues 199-203 | Kanamycin | pBR322 |
| pSML05 | T7-RNAP with DogTag insertion at residues 200-203 | Kanamycin | pBR322 |
| pSML06 | T7-RNAP with DogTag insertion at residues 276-278 (1) | Kanamycin | pBR322 |
| pSML07 | T7-RNAP with DogTag insertion at residues 276-278 (2) | Kanamycin | pBR322 |
| pSML08 | T7-RNAP with DogTag insertion at residues 355-362 | Kanamycin | pBR322 |
| pSML09 | T7-RNAP with DogTag insertion at residues 357-363 | Kanamycin | pBR322 |
| pSML10 | T7-RNAP with DogTag insertion at residues 468-472 | Kanamycin | pBR322 |
| pSML11 | T7-RNAP with DogTag insertion at residues 566-568 | Kanamycin | pBR322 |
| pSML12 | T7-RNAP with DogTag insertion at residues 587-592 | Kanamycin | pBR322 |
| pSML13 | T7-RNAP with DogTag insertion at residues 591-593 | Kanamycin | pBR322 |
| pSML14 | T7-RNAP with DogTag insertion at residues 708-712 | Kanamycin | pBR322 |
| pSML15 | T7-RNAP with DogTag insertion at residues 710-712 | Kanamycin | pBR322 |
| **T7-RNAP expression plasmids with manually designed insertions** | | | |
| pSML16 | T7-RNAP with DogTag insertion at residues 563-564 | Kanamycin | pBR322 |
| pSML17 | T7-RNAP with DogTag insertion at residues 599-600 | Kanamycin | pBR322 |
| pSML18 | T7-RNAP with DogTag insertion at residues 713-714 | Kanamycin | pBR322 |
| **T7-RNAP expression plasmids with SIMPLIFE multi-state designed single insertions** | | | |
| pSML19 | T7-RNAP with DogTag insertion at residue 97(1) | Kanamycin | pBR322 |
| pSML20 | T7-RNAP with DogTag insertion at residue 97(2) | Kanamycin | pBR322 |
| pSML21 | T7-RNAP with DogTag insertion at residue 130(1) | Kanamycin | pBR322 |
| pSML22 | T7-RNAP with DogTag insertion at residue 230(1) | Kanamycin | pBR322 |
| pSML23 | T7-RNAP with DogTag insertion at residue 230(2) | Kanamycin | pBR322 |
| pSML24 | T7-RNAP with DogTag insertion at residue 96 | Kanamycin | pBR322 |
| pSML25 | T7-RNAP with DogTag insertion at residue 203(1) | Kanamycin | pBR322 |
| pSML26 | T7-RNAP with DogTag insertion at residue 203(2) | Kanamycin | pBR322 |
| **T7-RNAP expression plasmids with SIMPLIFE multi-state designed multiple insertions** | | | |
| pSML27 | T7-RNAP with DogTag insertions at residues 3–5 and 357–363 | Kanamycin | pBR322 |
| pSML28 | T7-RNAP with DogTag insertions at residues 3–5 and 591–593 | Kanamycin | pBR322 |
| pSML29 | T7-RNAP with DogTag insertions at residues 357–363 and 591–593 | Kanamycin | pBR322 |
| **Reporter and mutational attachment expression plasmids** | | | |
| pRep01 | mCherry expressed by a T7 promoter | Gentamycin | p15A |
| pRep02 | DogCatcher expressed by a $P_{CymRC}$ promoter promoter | Kanamycin | p15A |
| pRep03 | mCherry expressed by a T7 promoter and DogCatcher::PmCDA1 expressed by a $P_{CymRC}$ promoter | Gentamycin | p15A |
| pRep04 | mCherry expressed by a T7 promoter and DogCatcher expressed by a $P_{CymRC}$ promoter | Gentamycin | p15A |

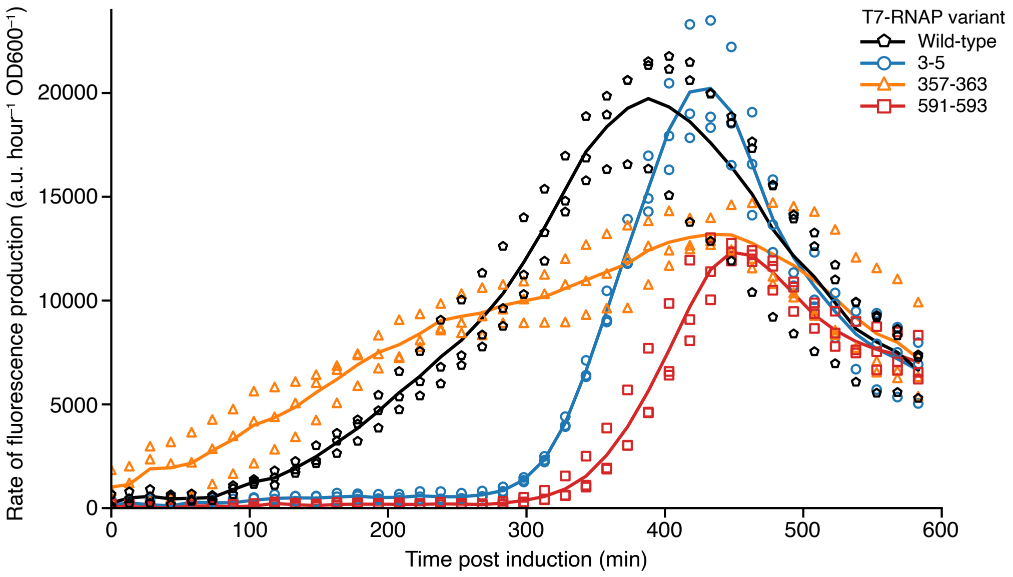

**Supplementary Figure 1: Varying fluorescence production over time for the modified T7-RNAPs.** mCherry fluorescence production rate (a.u./hour) per OD600 over time for wild-type T7-RNAP and three tagged T7-RNAPs at residues 3–5, 357–363 and 591–593. Maximum production rates peak at around mid-exponential phase (around 7 hours), but ordering of rates between T7-RNAP variants over time differs at earlier and later growth phases.

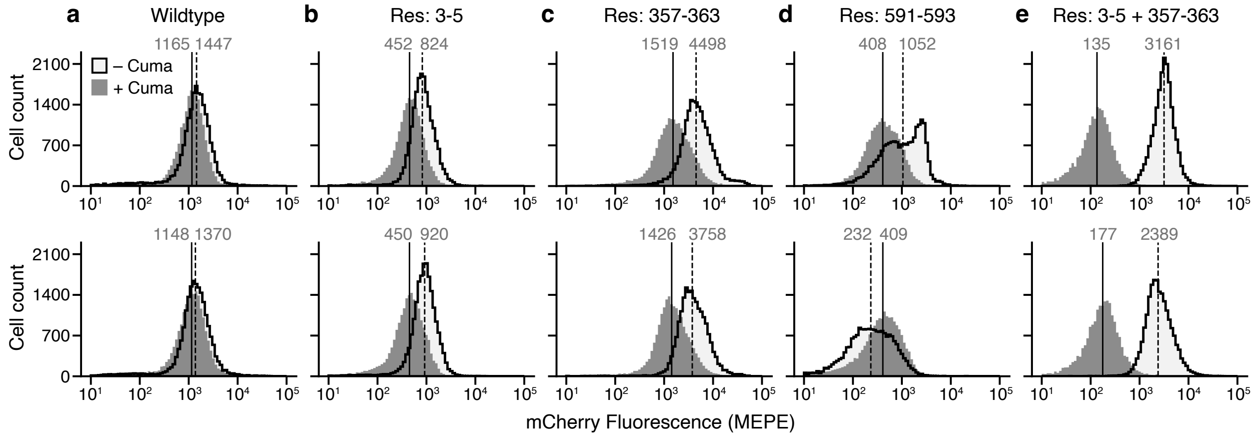

**Supplementary Figure 2: Impact of mutational attachments on mCherry fluorescence distributions.** Fluorescence distributions of mCherry from flow cytometery for different T7-RNAP variants after 32 hours, with (solid grey) and without induction (light grey and solid black outline) by cuminic acid. Data for biological replicates shown as columns. Solid and dashed vertical lines denote the medians for each distribution with cuminic acid present and absent, respectively, with their median values shown above each line in grey. Fluorescence measurements given in Molecules of Equivalent PE per cell (MEPE) units. Data shown for two biological replicates, third replicate can be found in **Figure 4b** of the main text. (**a**) Wild-type T7-RNAP. (**b**) T7-RNAP with an insertion at residues 3–5. (**c**) T7-RNAP with an insertion at residues 357–363. (**d**) T7-RNAP with an insertion at residues 591–593. (**e**) T7-RNAP with insertions at residues 3–5 and 357–363.
